# Pitch discrimination performance of ferrets and humans on a go/no-go task

**DOI:** 10.1101/165852

**Authors:** Kerry MM Walker, Amelia Davies, Jennifer K Bizley, Jan WH Schnupp, Andrew J King

## Abstract

Animal models are widely used to examine the neurophysiological basis of human pitch perception, and it is therefore important to understand the similarities and differences in pitch processing across species. Pitch discrimination performance is usually measured using two-alternative forced choice (2AFC) procedures in humans and go/no-go tasks in animals, potentially confounding human-to-animal comparisons. We have previously shown that pitch discrimination thresholds of ferrets on a 2AFC task are markedly poorer than those reported for go/no-go tasks in other non-human species (Walker *et al*., 2009). To better compare the pitch discrimination performance of ferret with other species, here we measure pitch change detection thresholds of ferrets and humans on a common, appetitive go/no-go task design. We found that ferrets’ pitch thresholds were ~10 times larger than that of humans on the go/no-go task, and were within the range of thresholds reported in other non-human species. Interestingly, ferrets’ thresholds were 100 times larger than human thresholds on a 2AFC pitch discrimination task using the same stimuli. These results emphasize that sensory discrimination thresholds can differ across tasks, particularly for non-human animals. Performance on our go/no-go task is likely to reflect different neurobiological processes than that on our 2AFC task, as the former required the subjects only to detect a pitch change while the latter required them to label the direction of the pitch change.

**ABBREVIATIONS:**

2AFC: 2-Alternative Forced Choice
F0: Fundamental Frequency

**HIGHLIGHTS:**

- Pitch discrimination thresholds of ferrets were 10 times larger than those of humans on a go/no-go task
- Ferrets’ pitch thresholds are similar to those reported for a range of other mammals
- Pitch thresholds of ferrets, but not humans, were drastically better on the go/no-go task than a 2AFC task using the same stimuli

## 1. INTRODUCTION

The brain makes sense of sound signals by extracting a limited number of perceptual features that are useful in identifying, localizing, and discriminating objects. Among these features, “pitch” is one of the most behaviorally relevant to humans and non-human animals. Pitch is defined by the American National Standards Institute (ANSI, 1994) as “that attribute of auditory sensation in terms of which sounds may be ordered on a scale extending from high to low”. The main acoustical correlate of pitch is the repetition rate of the sound waveform. Perceptually, it is the tonal quality of a sound that we use to form musical melodies, and which also plays a key role in verbal communication. The pitch of a voice is used to identify the prosody, gender (Lass *et al*., 1976) and emotional state of a speaker (Johnson, 1990), and allows us to effectively attend to the speech of one person in a crowded room (de Cheveigné *et al*., 1997). The perception of pitch is similarly utilized in communication among non-human animals, as many species interpret vocal calls (Hauser and Fowler, 1992; Miller and Hauser, 2004) and determine speaker identity (Capranica, 1966; Kojima *et al*., 2003) based on pitch cues.

The commonalities of pitch perception among humans and other species is important, as it allows us to examine the neurophysiological basis of this phenomenon in animal models. A number of research groups have investigated the representation of sound periodicity in the spiking responses of auditory cortical neurons using a variety of animal models (e.g., cat: Eggermont, 1991; macaque: Steinschneider *et al*., 1998; marmoset: Bendor and Wang, 2005; ferret: Bizley *et al*., 2010). However, the similarity between the pitch perception of some of the more common animal models of hearing and that of human listeners remains unclear. For example, recent studies have shown that marmosets and humans differ in their use of resolved harmonic pitch cues (Osmanski *et al*., 2013). Such psychophysical details are crucial to validating the relevance of findings in animal studies to human pitch perception, particularly because human pitch perception may be specialized through speech and music experience.

Although the aforementioned ANSI definition of pitch specifically refers to the ordering of sounds, most investigations of pitch discrimination in animals have used a go/no-go paradigm in which the subjects are required to detect a periodicity change but do not need to label that change as "upward" or "downward" along an ordered scale. In a go/no-go task, the subject is typically required to responds to a pitch deviant in an on-going sequence of sounds in order to obtain reward, (commonly, water or food), or to avoid a punishment, (commonly, a mild electrical shock). Previous studies that have trained animals to detect a change in the pitch of a 500-Hz pure tone on a go/no-go task have found the smallest detectable frequency difference to be between 7 – 18 Hz for a number of mammalian species, including chinchillas (Nelson and Kiester, 1978), cats (Elliott *et al*., 1960), guinea pigs (Heffner *et al*., 1971) and tree shrews (Heffner *et al*., 1969). Monkeys tend to have better frequency acuity than the above species, with 500-Hz frequency difference thresholds of 4 – 5.5 Hz (Massopust *et al*., 1971; Stebbins, 1973). Fewer studies have measured animals’ pitch discrimination thresholds using complex sounds.

In a recent study, we used a 2-Alternative Forced Choice (2AFC) task to measure ferrets’ pitch discrimination thresholds, and we used both pure tones and artificial vowels as stimuli (Walker *et al*., 2009). On each trial of our task, ferrets were presented with two consecutive sounds and were required to indicate whether the pitch of the second sound was lower or higher than that of the first. This task was methodologically similar to the 2AFC task commonly used to measure human pitch discrimination thresholds, in which two sounds are presented and the subject is asked to report whether the first or second sound was higher in pitch (e.g. Moore, 1973; Wier *et al*., 1977; Won *et al*., 2010). Ferrets produced similar pitch thresholds for pure tones and artificial vowels, suggesting that the pitch discrimination of simple and complex sounds can be compared across studies. The average pitch Weber fraction across ferrets was found to be 36.4%, corresponding to a pitch difference threshold of 182 Hz for a 500 Hz reference (Walker *et al*., 2009). This is an order of magnitude higher than the thresholds reported above for other species, which may reflect: a) genuinely poorer pitch acuity in ferrets compared to other mammals, or b) the effects of different task demands in the go/no-go and 2AFC paradigms.

Comparative psychoacoustic studies have usually related human pitch discrimination measured on 2AFC tasks to the thresholds of animals on go/no-go tasks (Fay, 1988; Shofner, 2000). If listeners’ psychophysical thresholds differ across go/no-go and 2AFC tasks, then this across-task confound may be problematic. Here, we test the pitch discrimination performance of humans and ferrets on a go/no-go task in order to aid comparison with previous studies in other species. By comparing the results of these experiments with those of our previous 2AFC task, we are also able to investigate whether task design affects the pitch difference thresholds of humans and ferrets.

## 2. MATERIALS AND METHODS

### 2.1 Subjects

Eight adult pigmented ferrets (1 male) were used in this study, which were housed individually or in pairs, with *ad lib* access to food. They were tested twice daily, and typically completed between 60-120 trials per session. On training days, the ferrets’ daily water consumption was limited to 60 ml/kg body weight. They were given water *ad lib* on non-experimental days. Regular otoscopic examinations and typanometry were carried out to ensure that both ears of the animals were clean and healthy. All experimental procedures were approved by the relevant ethical review committees at the University of Oxford, and the animal studies were carried out under licence from the UK Home Office in accordance with the Animals (Scientific Procedures) Act 1986.

### 2.2 Ferret training apparatus

The apparatus used for training has been described in a previous publication (Walker *et al*., 2009). Sound-insulating chambers were made of double-glazed walls and floors, with solid wood ceilings (Fig. 1A). Each chamber contained a plexiglass back wall, and a row of three stainless steel waterspouts protruded through this wall. For some animals, the licking response at the waterspout was detected as a small change in electrical current between the spout itself and a metal plate on which the ferret stood (Hayar *et al*., 2006). For the remaining animals, each waterspout was located inside a cylindrical “nose-poke hole” with an infrared light-emitting diode and photodetector pair at its entrance. The ferret’s response at the waterspout was detected when the infrared beam was interrupted by the ferret’s snout. We observed that ferrets could trigger either of these response systems reliably and with similar ease. All sound stimuli were presented from a loud speaker positioned above the center spout (Visaton FRS 8; flat response within ±2 dB from 200 - 20,000 Hz). In all behavioral experiments in this study, the sound presentation, behavioral measurements, and water delivery were controlled by custom MATLAB (The MathWorks, Inc.) scripts running on a personal computer, and a TDT RM1 signal processor (Tucker-Davis Technologies, Alachua, FL).

**FIG. 1.**
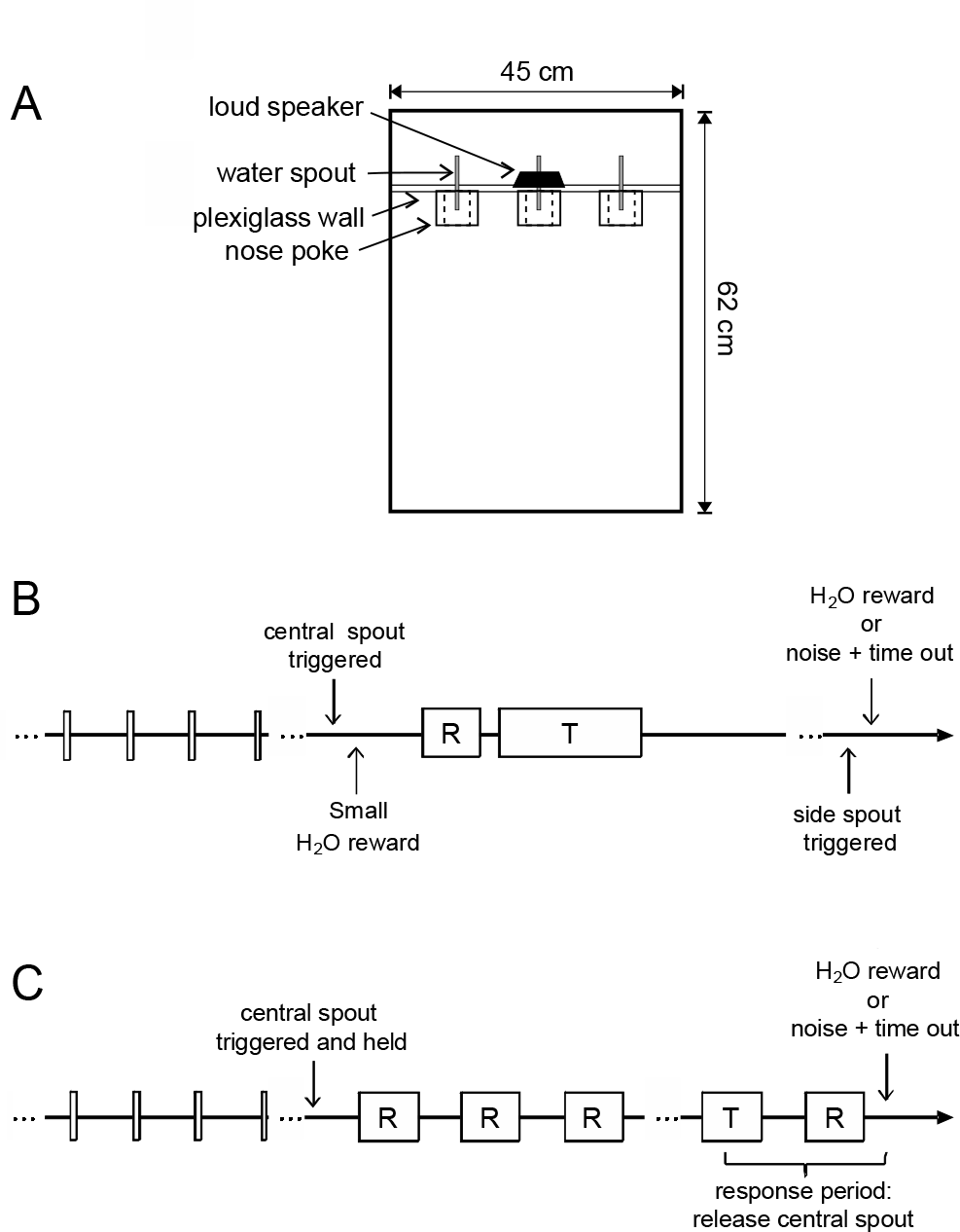
Ferret testing apparatus and task design. **A** Ferret testing chamber, viewed from above. Ferrets’ responses were detected by infrared beams across each of the three nose-poke holes, and water rewards were delivered through the water spouts situated inside these holes. **B** Trial schematic for the go/no-go task. Rectangles positioned along the time line represent the timing of auditory stimuli: the ready signal (a short, repeating pure tone), followed by a variable number of presentations of the artificial vowel at the reference F0 (“R”), a presentation of the vowel at a target F0 (“T”), and a final presentation of the reference F0 (“R”). **C** Trial schematic for the 2AFC task. The timing of the ready signal, and vowel at the reference (“R”) and target (“T”) F0 are shown as in B.

### 2.3 Artificial vowel stimuli

The sound stimuli used in these experiments were artificial vowel sounds, consisting of band-pass filtered click trains. The repetition rate of the click train determined the fundamental frequency (F0) of the sound, which is a strong acoustical determinant of perceived pitch. To make these sounds more naturalistic, they were band-passed with filters centered on 430, 2132, 3070 and 4100 Hz. These pass bands correspond to the first four formants of the English vowel /*i*/. It was not necessary to use sounds that model human speech to answer our current experimental question, but this stimulus was chosen to facilitate comparisons with our earlier work. Stimuli were cosine ramped on and off (5 ms duration), and were pseudorandomly varied in intensity across a 15 dB range from 65-80 dB SPL in all behavioral tasks to discourage the use of intensity cues.

### 2.4 Go/no-go pitch change detection task

Five naïve ferrets (namely, F4, F5, F6, F7 and F8) were trained to perform a positively reinforced go/no-go pitch change detection task. All 5 animals received 1-3 months of training prior to experimental testing, depending on the performance of the animal on several training stages. Training stages were modeled after those used successfully in previous studies (Prosen *et al*., 1989; Talwar and Gerstein, 1998). Ferrets were first trained to nose-poke at the central spout for a water reward, which was always delivered from the right waterspout. The time required to remain at the central spout in order to receive a water reward was gradually lengthened across trials and sessions, until the ferrets learned to nose-poke continuously for 2 s.

A “target” vowel (F0 = 600 Hz) presentation was next introduced. In order to receive a reward, the ferrets were required to remain at the central nose-poke hole until the target was presented. *False Alarms* (i.e., moving from the central spout before the target onset) resulted in the presentation of a broadband noise burst (500 ms duration, 70-75 dB) and a time-out of 12 s. A repeating 5-kHz pure tone (20 ms duration, 0.5 ms rise/fall time, and 200 ms inter-tone interval) was presented following each reward or time-out to indicate to the animal that the central spout could now be triggered to initiate the next trial.

Once ferrets achieved >85% correct on this task for 2 sessions in a row, we introduced a repeating “reference” sound into the start of the trial. The reference was the same vowel as the target, but had an F0 of 400 Hz. All vowels (references and targets) were separated by 200-ms periods of silence. Each trial consisted of a sequence of reference sounds at 400 Hz, followed by a target at 600 Hz, and then a final 400-Hz reference (see Fig. 1B). The number of reference sounds presented prior to target onset varied randomly across trials. For three animals, the number of reference sounds preceding the target ranged between 2-6, and the duration of each vowel was 250 ms. For the final two animals, 2-3 references were presented before the target on each trial, and the sounds were each 350 ms in duration.

The response window began at the onset of the target, lasted throughout the duration of the target and following reference, and terminated at the end of the silent interval following the final reference. Movements from the central spout during this window were considered *Hits* and resulted in a water reward. Center spout releases occurring before the target onset were considered *False Alarms*, and were negatively reinforced with a broadband noise and 12 s time-out, as above. If the ferret failed to respond by the end of the response window, it was considered to have *Missed* the target, and this resulted in a broadband noise burst followed by a 4 second time-out.

The reference was first introduced at an average level 40 dB below the target, so that the animal was required to respond to the onset of the target in a relatively quiet background. This level difference was gradually diminished across sessions by increasing the reference level 5-10 dB after each session in which the ferret achieved a score of >85%. Each animal reached the final stage of behavioral testing when the average intensity levels of the reference and target were equal. When the ferret completed two such consecutive sessions at >85% correct, the ferret began experimental testing.

In the testing stage, the F0 of the target was varied across trials. In some sessions, the target F0 took values higher than the reference, and, in separate sessions, the target was presented at F0’s below the 400-Hz reference. A randomly interspersed 10% of trials were “catch trials”, which consisted of a sequence of reference sounds without a target. *Correct Rejections* on such trials, in which the ferret kept licking the central spout until the end of the sequence, were rewarded with water.

### 2.5 2AFC pitch direction judgment task

Five ferrets (1 male) were trained on a series of training tasks in order to shape their behavior for a 2AFC pitch discrimination task. Three of these animals (namely, F1, F2 and F3) were naïve to training, and their performance on the 2AFC task has been previously reported in detail (Walker *et al*., 2009). Here, we report on only the subset of their testing sessions in which the F0 of the reference was 400 Hz (±10 Hz). The 2 other animals (namely, F4 and F5) trained on this task were first trained on the go/no-go task described below. The same apparatus and stimuli were used for the go/no-go and 2AFC tasks.

Details of 2AFC training and testing are available in Walker *et al*., 2009, and so only the experimental testing is described in brief here. A trial schematic of the final testing stage of the 2AFC task is shown in Fig. 1C. At the beginning of each trial, a repeating tone burst (5 kHz tone, 20 ms duration, 0.5 ms rise/fall time, and 200 ms inter-tone interval) was presented, indicating to the animal that it could now trigger the presentation of the vowel stimuli by nose poking at the central response spout. The center nose poking was reinforced on 10% of trials with a small water reward (0.1-0.2 ml). Two artificial vowel sounds were presented following the center nose-poke: a 400 Hz reference vowel (200 ms duration), followed by a target vowel (500 ms duration) that varied in F0 from trial-to-trial. The ferret was required to respond to the target vowel at one of the two peripheral nose poke holes, depending on the relative F0 of the target and reference. If the target was higher than the reference, right spout choices were rewarded with water (0.3-0.5 ml per trial). If the F0 decreased from reference to target, a left spout choice resulted in a water reward. Incorrect spout choices resulted in the presentation of a broadband noise burst (500 ms duration, 70-75 dB SPL), followed by a time-out of 10-12 s. Trials were reset without time-out if the animal made no response within 15 s of target offset.

### 2.6 Human psychophysical testing

The pitch discrimination thresholds of 5 adult humans (3 female, 2 male, ages 19-31) were measured on go/no-go and 2AFC tasks designed to be very similar to those executed by the ferrets. The task order was counterbalanced across subjects. Testing took place inside a sound-attenuated booth (Industrial Acoustics Company, Winchester, UK) and stimuli were presented diotically through headphones (HD 25-1; Sennheiser UK Ltd, High Wycombe, UK). Subjects made responses by pressing keys on a computer keyboard, and feedback was given visually on a computer monitor after each trial (“correct” or “incorrect”). Otherwise, the two human psychophysical tasks were procedurally identical to those used to test ferrets’ pitch thresholds. The same artificial vowel sounds were used as stimuli in all tasks, and these were presented with the same level randomization (65-80 dB SPL) across trials as used in the ferrets’ tasks. In the human go/no-go task, vowels were 250 ms in duration and 2-6 references preceded the target on each trial.

Each task was explained to the subject by the experimenter, and the subject was then given a limited number (n ≈ 25) of practice trials to ascertain that they understood the procedure before psychophysical testing commenced. On the go/no-go task, the subject was asked to respond as quickly as possible without making errors to a change in the pitch of the repeating sound. For the 2AFC task, they were instructed to indicate whether the target sound was higher or lower in pitch than the reference.

Humans perform much better than ferrets on pitch discrimination tasks, so the pitch differences presented to humans were smaller than those required to estimate ferrets’ difference thresholds. The exact pitch differences presented to each subject were chosen based on each individual’s performance on practice trials prior to testing. In order to match the task design used for ferrets, we did not use an adaptive procedure but instead chose values based upon subjects’ performance in the first 50-100 trials.

### 2.7 Data analysis

Data were pooled across behavioral sessions for each subject for the purposes of analysis. Chance performance on the 2AFC task was 50%. Chance performance on the go/no-go task was generally lower than this, but is more difficult to calculate as it depends on both the animal’s strategy (i.e. false alarm rate as a function of position in the stimulus sequence) and the number of possible target positions. If we assume that the animals randomly responded with equal probability to one of the possible target positions in this sequence on each trial, chance performance would be 30% when there are 3 possible target positions and 18% when the target has 5 possible positions (taking into account the 10% chance of a catch trial). Our analyses excluded 2AFC sessions in which the ferret performed <60% of trials correctly, and go/no-go sessions in which either <40% of trials were performed correctly or the false alarm rate was >65%. These exclusion criteria led us to discard a small number of sessions in which well-trained animals performed the task poorly overall.

On the 2AFC task, responses at each pitch difference were expressed as a percentage of “pitch increase” judgments (i.e. right spout choices). These observed values were then fitted, using generalized linear model regression for binomial distributions, to a cumulative logistic distribution function. The F0 difference threshold for the 2AFC task was calculated as the F0 difference required for the subject to achieve 76.02% correct performance, which corresponds to a discriminability index (*d*’) value of 1 on the equivalent change detection task (Wickens, 2002).

For the go/no-go task, the hit rate for each target F0 condition was calculated as the proportion of trials on which the subject correctly responded to the target in the appropriate response window (i.e., the *Hit rate*). The *False Alarm rate* was calculated as the proportion of erroneous responses on catch trials. Discriminability of each target can be expressed as *d’*, calculated as:

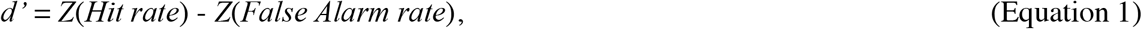

where *Z* is the inverse of the cumulative Gaussian distribution (Green and Swets, 1974).

Pitch difference thresholds for the go/no-go task were initially calculated separately for sessions in which the F0 of the target increased compared the reference, and those in which the F0 decreased.

Discriminability performance for pitch increases was calculated as *d*’ using Eq.1, and that for pitch decreases was represented as –*d*’, resulting in a sigmoidal discriminability curve across the full target F0 range. These raw values were then fitted using Maximum Likelihood Estimation to the following logistic function:

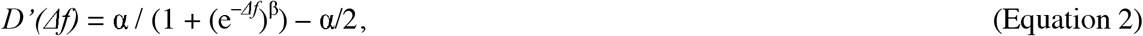

where Δ*f* is the F0 difference between the target and reference, α is a constant that rescales and shifts the function in the y-direction, and β is a constant that rescales the function in the x-direction. An estimate of the F0 difference threshold for the go/no-go task was then calculated as the point on this fitted function where *D’* = 1.

Pitch difference thresholds were compared across tasks within the same subjects using paired Mann-Whitney U tests (*U*), and across different subject groups using Wilcoxon Rank Sum tests (*T*). An alpha of 0.05 was used throughout as criterion for statistical significance.

## 3. RESULTS

### 3.1 Pitch change detection in ferrets

Five naïve ferrets were trained to detect an F0 change in a repeating artificial vowel sound. On each trial, the ferret initiated the vowel sequence by nose-poking at the center spout and was required to remain in the nose-poke hole until an F0 change occurred (Fig. 1C). Hits and Correct Rejections were rewarded with water, whereas False Alarms and Misses resulted in a time-out. Both Hit and False Alarm rates increased throughout the sound sequence (Fig. 3A). These response rates were relatively consistent across individual animals (Fig. 3A; symbols), but were dependent upon the maximum length of sound sequence presented to the animal. For example, the three animals that were trained with up to 8 sequential sounds per trial (Fig. 3A; circles, diamonds and squares) false alarmed to the fourth sound in the sequence on less than 15% of trials, and the average hit rate for targets at position 4 in the sequence was near 80%. For the other two animals, trials consisted of no more than 5 sounds in the sequence, and therefore there was a greater probability of targets occurring in the third or fourth sound in the sequence. Accordingly, these ferrets false alarmed to the fourth sound in the sequence at a rate of near 40%, while their hit rate for targets in this position was over 90% (Fig. 3A; triangles and inverted triangles). These results suggest that ferrets adapted their response rate to the task statistics in order to improve their overall performance.

**FIG. 2.**
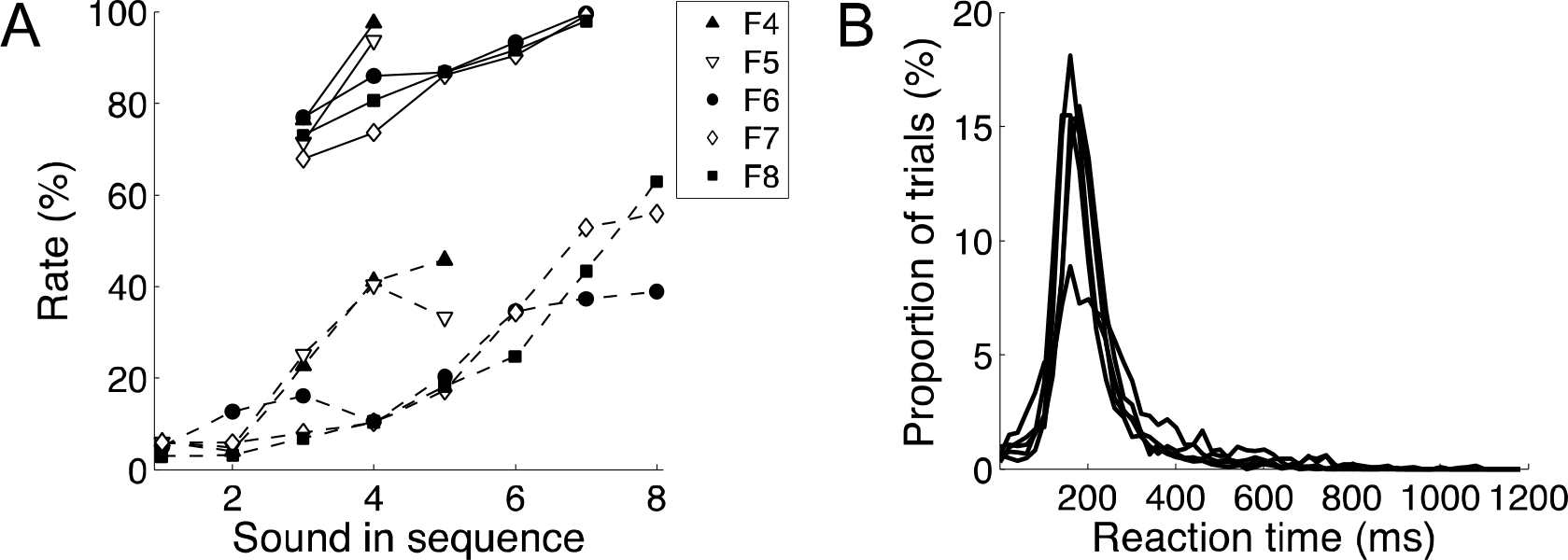
Ferret performance statistics for the go/no-go task. **A** Hit rates (solid lines) and FalseAlarm rates (dashed lines) are plotted as a function of the position in the stimulus sequence at which the response occurred. Data from individual ferrets are distinguished by different symbols (see legend). **B** Distribution of reaction times with respect to target onset, for trials in which the ferret correctly detected the pitch change (i.e. Hits). Each line represents an individual ferret, and the outlier with the smallest peak at 200ms is F4.

In our go/no-go task, ferrets could correctly respond to the pitch change any time following the onset of the target sound through to the end of the following reference sound. They were thus permitted to respond to either the initial pitch change (from the reference to target F0) or the second pitch change (from target F0 back to reference F0). Reaction times were calculated as the time at which the animal withdrew its head from the central nose-poke hole. The distribution of reaction times with respect to target onset show that on the vast majority of “hit” trials, ferrets responded during the target presentation, and almost always before the onset of the final reference sound (Fig. 3B). The mean (± SEM) reaction time across all animals was 227.2 ± 9.0 ms.

F0 change detection thresholds were determined for the go/no-go task based on a psychometric fit to *d*’ values for each ferret (Fig. 4). The difference threshold was estimated from this fitted curve at the point *D’* = 1. *D*’ values for F0 increases (Fig. 4; circles) and decreases (Fig. 4; crosses) were measured in separate behavioral sessions, and so pitch thresholds were initially estimated separately from each of these two datasets. In these analyses, pitch change detection was found not to differ significantly between F0 increases and decreases (Fig. 5A; *T* = 3, *p* = 0.313). Therefore, for all subsequent analyses, an overall F0 difference threshold was calculated for each animal by fitting the entire go/no-go dataset to the logistic function in Eq. 2, as shown in Fig. 4.

**FIG. 3.**
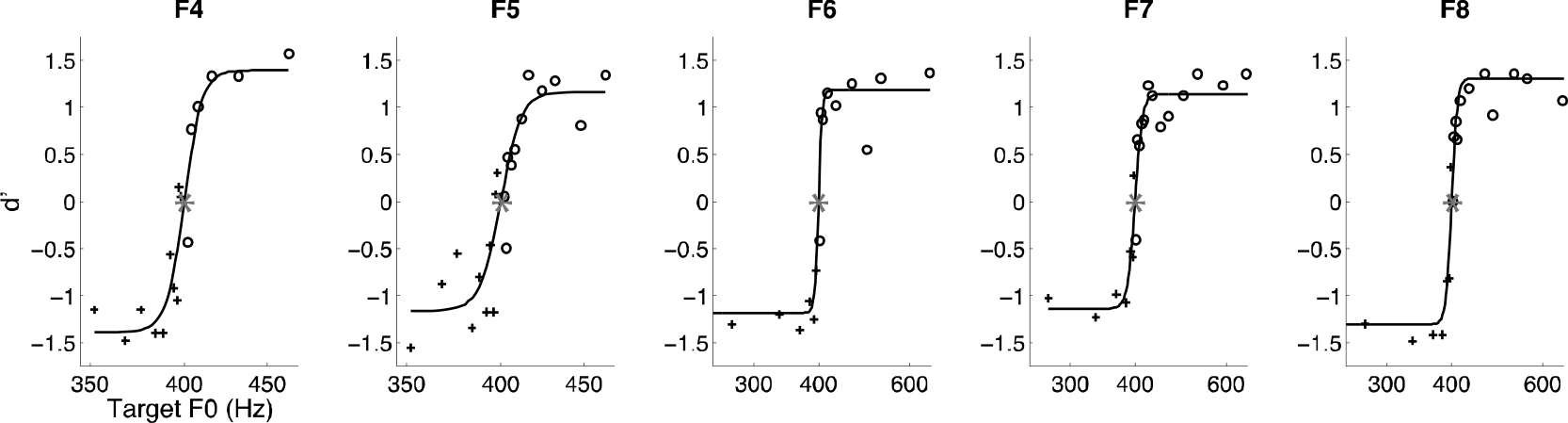
Ferret pitch discrimination performance on the go/no-go task. Each graph shows the psychophysical performance of one of the five ferrets tested. For each ferret, raw *d’* scores are plotted a function of the target F0 (symbols). The psychometric curves (Eq. 2) fitted to these data are shown (lines). Performance during F0 increase (circles) and F0 decrease (crosses) are plotted separately. Performance on F0 decreases are expressed as –*d*’ for the purposes of curve fitting. The reference F0 (400 Hz) is indicated with an asterisk at *d’*=0. The ferret name is indicated at the top of each graph. Note that different target ranges are shown on the x-axes for ferrets F4 and F5, compared to F6, F7 and F8.

### 3.2 Pitch direction judgments in ferrets

Five ferrets were trained to discriminate pitch increases from decreases on a 2AFC task. On each trial, an artificial vowel sound was presented at a set “reference” F0, followed by a “target” vowel that varied in F0 across trials (Fig. 1B). Ferrets were required to respond to the target by licking the spout to their left if the target vowel was lower in periodicity than the reference, or a spout to their right if the target F0 was higher than the reference. The resulting psychometric curves of the 5 ferrets are shown in Fig. 2. For each ferret, the 76.02% pitch difference threshold was estimated as the average of the F0 differences corresponding to 76.02% and 23.98% right responses on the psychometric curve in Fig. 2. The pitch difference was found to be 131.5 Hz for F1, 192.0 Hz for F2, 346.5 Hz for F3, 294.5 Hz for F4, and 493.5 Hz for F5 (mean = 291.6 Hz; SEM = 63.0).

**FIG. 4.**
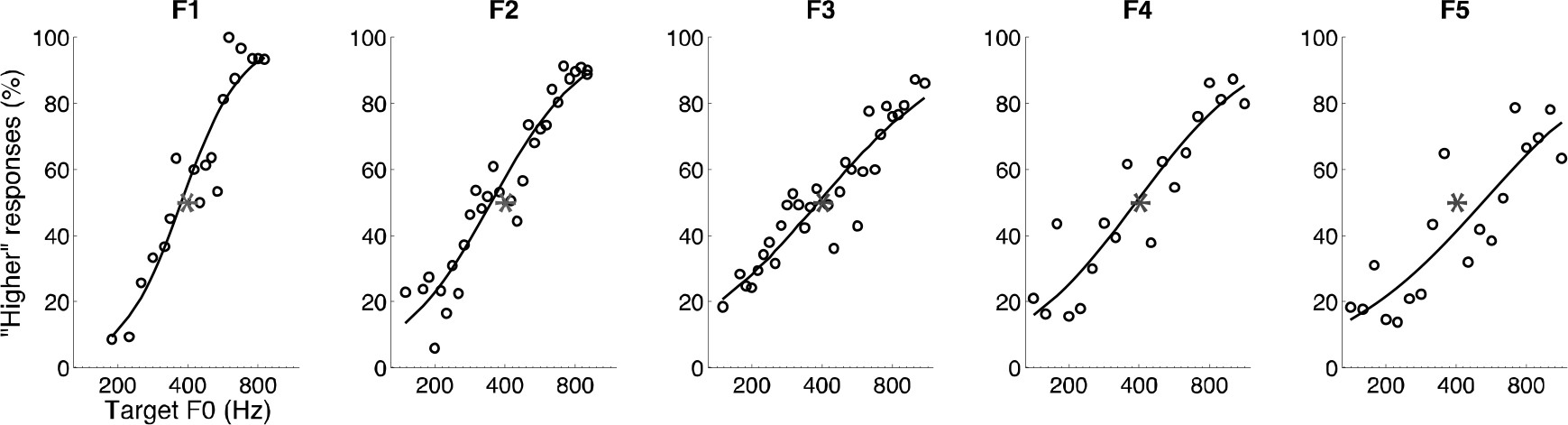
Ferret pitch discrimination performance on the 2AFC task. Each graph shows the psychophysical performance of one of the five ferrets tested (indicated by alphanumeric names, above). The proportion of right (“higher”) spout choices are plotted (circles) as a function of the target F0, and an asterisk indicates the reference F0 (400 Hz). Psychometric curves are fitted as cumulative Gaussian distributions (black lines).

### 3.3 Comparison of 2AFC and go/no-go performance in ferrets

The F0 change detection threshold was 14.4 ± 4.1 Hz (mean ± SEM) on the go/no-go task, which is over an order of magnitude better than the pitch difference threshold measured on the 2AFC task (Fig. 5B; *U* = 15, *p* = 0.008). We examined if this difference in thresholds on the two tasks might correspond to a large difference in the difficulty of learning the procedural aspects of each task. The mean (+ SEM) number of trials and sessions required by previously untrained animals to reach criterion on the final training stage are plotted in Fig. 5C and 5D, respectively. In both tasks, criterion was set as performing the final training stage at >85% correct on two consecutive sessions. Data from 5 naïve animals are shown as black symbols^1^. For comparison, the number of sessions and trials required by the two experienced animals to learn the 2AFC task following go/no-go training are indicated with gray symbols, but these values were not included in the group average. Note that the 2 experienced ferrets learned the 2AFC task more quickly than the other 5 animals, suggesting that some of the procedural learning they acquired on the go/no-go task (e.g. how to nose-poke at spouts to trigger sounds and receive rewards) transferred to the 2AFC task.

We did not find a significant difference between the two tasks in either the number of trials (*T* = 33, *p* = 0.310) or number of sessions (*T* = 33.5, *p* = 0.238) that naïve ferrets required to reach criterion. Therefore, our data suggest that the pitch threshold differences are unlikely to be due to differences in the acquisition demands of the two tasks.

**FIG. 5.**
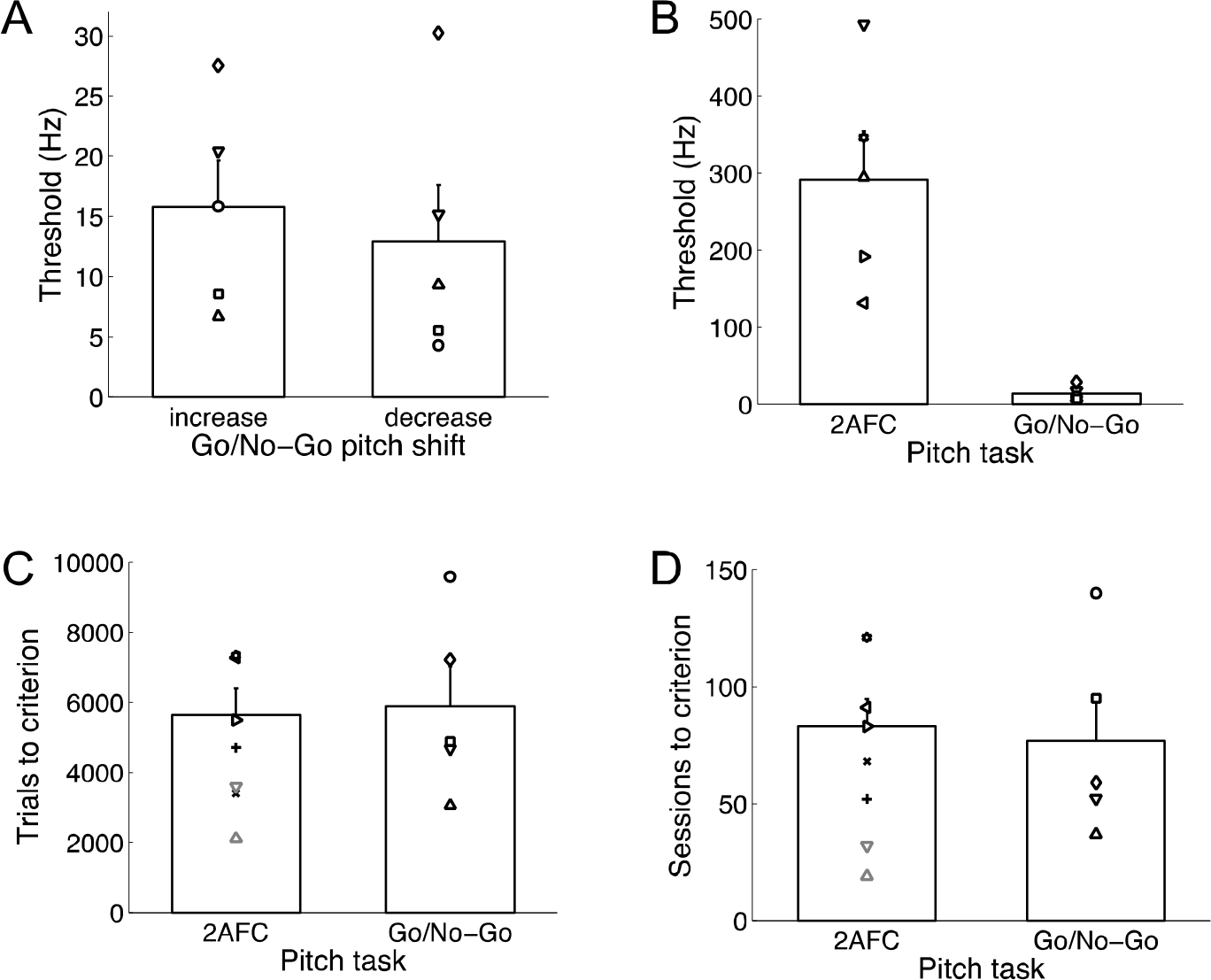
Ferret pitch discrimination performance indices compared between two tasks. Each plot shows the mean values plus SEM. The thresholds for individual ferrets are indicated with symbols. **A** The mean pitch difference thresholds of 5 ferrets on the pitch increase (left) and pitch decrease (right) versions of the go/no-go task. **B** Ferrets’ pitch difference thresholds measured on the 2AFC (left) and go/no-go (right) task. **C** Total number of pre-training trials performed before ferrets reached criterion on the final training stage of the 2AFC (left) and go/no-go (right) procedure. The 2 animals indicated in grey were re-trained on the 2AFC task following go/no-go training and so their scores are not included in the group average. **D** Number of pre-training sessions performed by ferrets prior to reaching criterion on the final stage of testing for the 2AFC (left) and go/no-go (right) procedure.

### 3.4 Comparison of human and ferret performance on 2AFC and go/no-go pitch discrimination tasks

Five human listeners were tested on button-press versions of the go/no-go and 2AFC pitch discrimination tasks, in which sound stimulation was presented over headphones. Go/no-go tasks of this form are highly uncommon in human psychophysics (with the exception of infant studies), so the False Alarm rates and reaction times of human subjects are provided for comparison to ferrets in Fig. 6A and B, respectively. Comparison with Fig. 3A shows that the False Alarm rates of humans were much lower than in the ferrets, while the Hit rates were also generally lower. Therefore, ferrets are more likely to “go” when unsure of a pitch change, whereas the human strategy appears to adhere to a higher criterion. This is consistent with the human reaction times (419.6 ± 14.8 ms, mean ± SEM; measured as keyboard button presses), which were almost twice as long as those of ferrets. Although their reaction times were longer, human subjects nevertheless usually responded to pitch changes before the onset of the final reference sound. Also in agreement with the ferret data, human thresholds on the go/no-go task did not differ significantly between sessions in which F0 increased (2.4 ± 0.3 Hz) and those in which it decreased (2.9 ± 0.9 Hz; *T* = 4, *p* = 0.438).

**FIG. 6.**
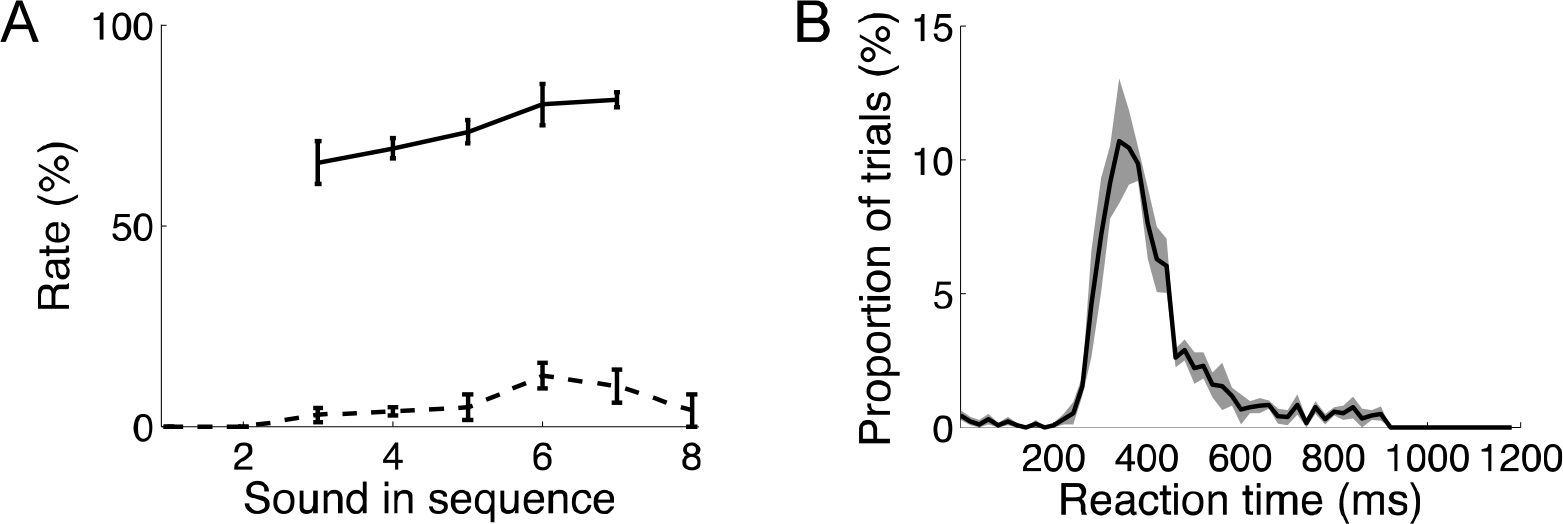
Human performance statistics relating to the go/no-go task. **A** Mean (± SEM) Hit rates (solid lines) and False Alarm rates (dashed lines) are plotted as a function of the position in the stimulus sequence at which the response occurred. **B** Mean (± SEM in grey) reaction times, with respect to target onset, on Hit trials.

Raw data and fitted psychometric curves of human listeners on the 2AFC and go/no-go tasks are shown in the upper and lower panels of Fig. 7, respectively. The same psychometric fitting procedures were used to derive difference thresholds from the ferret and human data, so that these values could be directly compared. In contrast to ferrets, which performed much better on the go/no-go than the 2AFC paradigm, the F0 thresholds of human listeners were found not to differ significantly between the 2AFC (1.1 ± 0.2 Hz) and go/no-go tasks (2.5 ± 0.4 Hz; *T* = 0, *p* = 0.063).

**FIG. 7.**
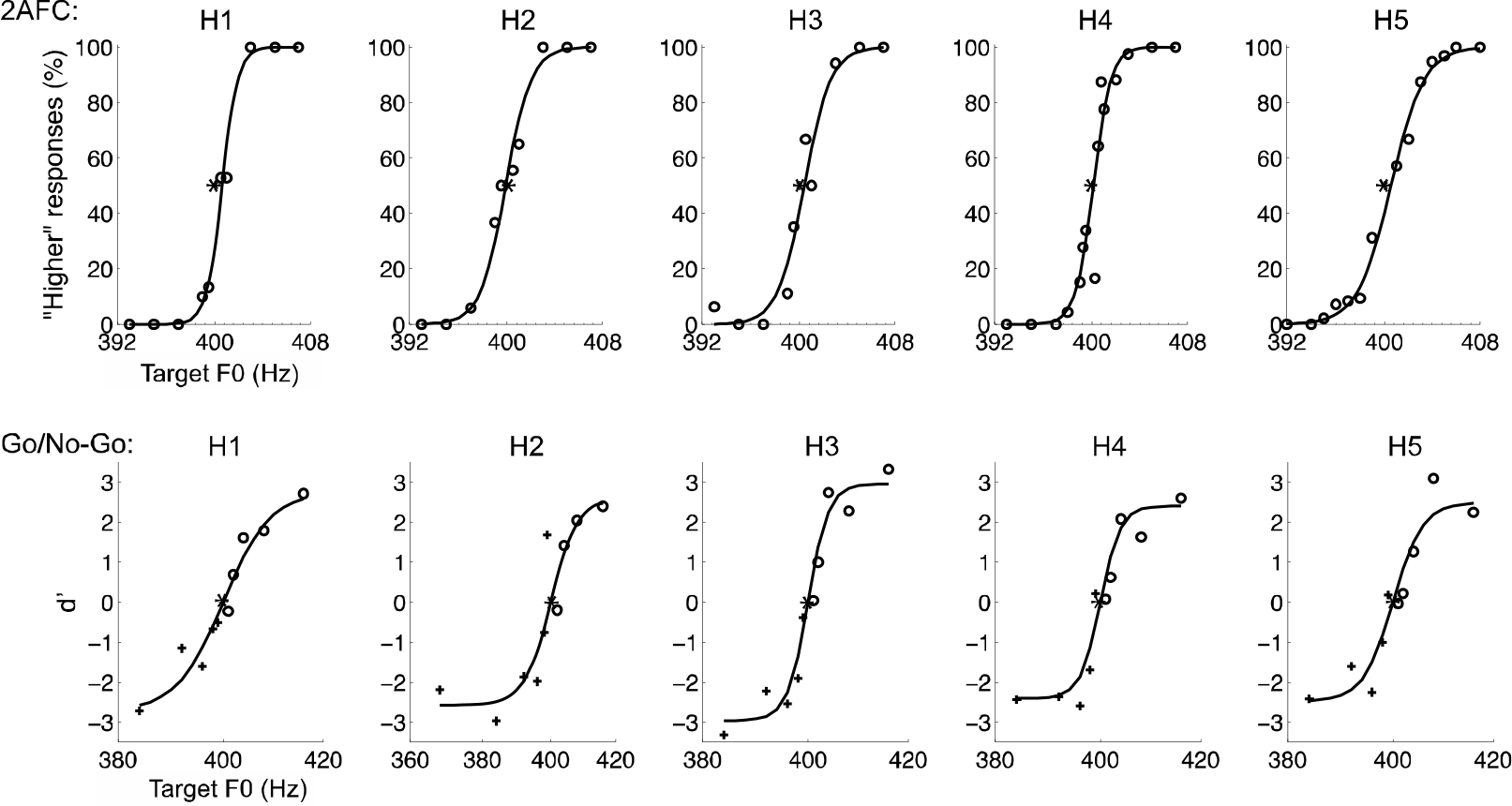
Pitch discrimination performance of human subjects on two tasks. Top row: Each graph shows the psychometric performance (circles) and fitted psychometric curve (line) of a different subject on the 2AFC task. Bottom row: Each graph shows the psychometric performance of a human listener on the go/no-go task, expressed as *d*’ for pitch increases (circles) and –*d*’ for pitch decreases (crosses). The fitted psychometric curve (line) is also shown. In all graphs, an asterisk at chance on the y-axis shows the F0 of the reference (400 Hz), and the subject code is indicated at the top of the plot.

In Fig. 8, the thresholds of ferret and human listeners are plotted side-by-side. Note that a log-Hz scale is used. Humans’ pitch thresholds were much better than those of ferrets on the 2AFC pitch direction judgment task (*U* = 40, *p* = 0.008). In fact, the average threshold of ferrets on this task was more than two orders of magnitude larger than that of human listeners (291.6 versus 1.1 Hz, respectively). Although the absolute difference in ferret and human mean thresholds on the go/no-go task (14.4 versus 2.2 Hz, respectively) was much smaller than on the 2AFC task, human listeners again performed significantly better than ferrets on the go/no-go task (*U* = 40, *p* = 0.008). The comparison commonly drawn by previous studies is that of human performance on a 2AFC pitch discrimination task and animals’ performance on a comparable go/no-go version of the task. By this comparison, we again found that the pitch discrimination of our human listeners was better than that of ferrets (*U* = 40; *p* < 0.008), with average thresholds of ferrets and humans being approximately an order of magnitude apart (14.4 versus 1.1 Hz, respectively).

**FIG. 8.**
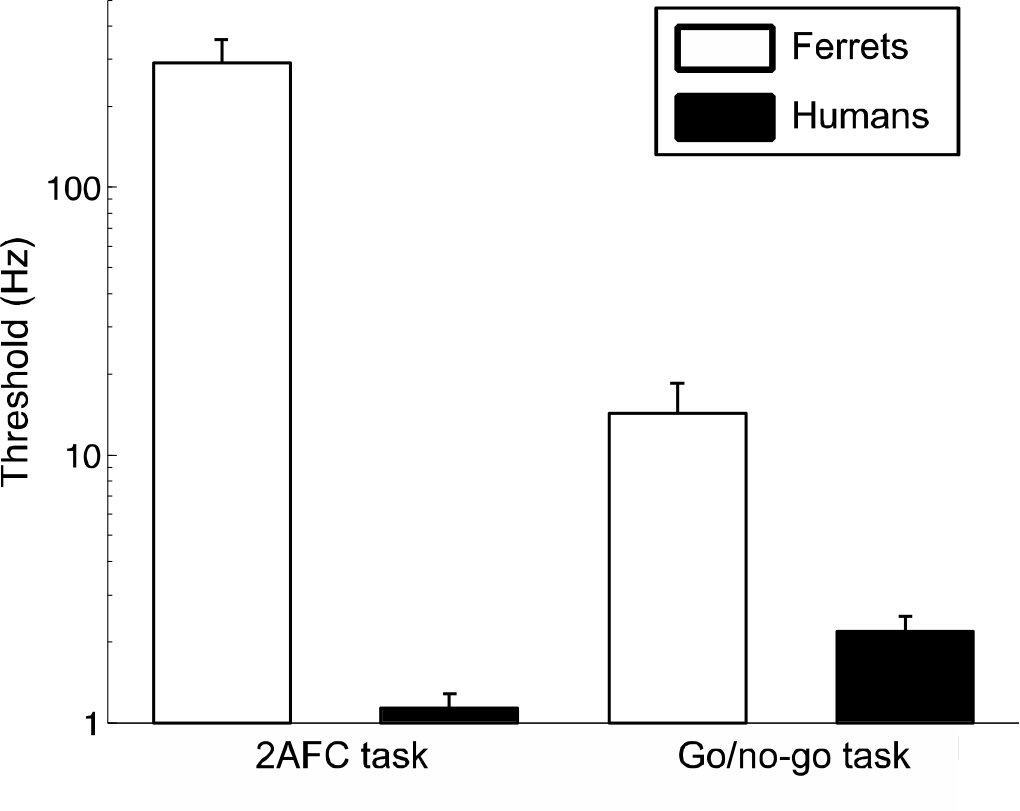
Comparison of pitch discrimination performance across two tasks and two species. The mean (+ SEM) pitch difference thresholds for ferret (white bars) and human (black bars) listeners on the 2AFC (left) and go/no-go (right) tasks are shown. Note that the y-axis is a logarithmic scale.

## 4. DISCUSSION

### 4.1 Comparisons of pitch discrimination performance between two tasks and two species

The majority of behavioral investigations of sound discrimination in animals have used go/no-go change-detection paradigms (Elliott *et al*., 1960; Heffner *et al*., 1969; Heffner *et al*., 1971; Massopust *et al*., 1971; Stebbins, 1973; Nelson and Kiester, 1978; Prosen *et al*., 1989), and so it has previously been unclear whether estimates of perceptual acuity (e.g. pitch difference thresholds) would depend on the type of task used. The present study demonstrates that the choice of behavioral task does, in fact, greatly affect the acuity threshold measured; ferrets’ pitch difference thresholds on a 2AFC high/low identification task are up to 10 times higher than their thresholds on a go/no-go change detection task in which the same stimuli are presented.

Psychophysical studies in adult human subjects commonly employ a 2AFC task in which two consecutive sounds are presented on each trial and the subject is asked to report whether the first or second sound is higher in pitch (Moore, 1973; Wier *et al*., 1977; Won *et al*., 2010). Such tasks have several advantages: they are trivial for human subjects to learn; each trial is quick to perform; they provide easily derivable difference thresholds; and, importantly, they are free from criterion bias (that is, an individual’s likelihood to report that an event occurred under uncertainty). Even pitch change detection studies in humans have often favored AFC tasks in which the listener reports which of two sequential pairs of tones contains a frequency change, known as the 4-interval AFC paradigm (e.g., Semal and Demany, 2006). Comparisons between the pitch discrimination thresholds of human and animal subjects have therefore often been made across task types, by comparing human thresholds on AFC tasks with those of animals on go/no-go tasks (Elliott *et al*., 1960; Fay, 1988; Shofner, 2000), without knowledge of how thresholds might vary across these two tasks within species.

We found that human F0 difference thresholds did not differ significantly between our 2AFC and go/no-go pitch discrimination tasks. Sinnott *et al*. (1992) and Klinge and Klump (2009) also found that humans’ thresholds on a go/no-go version of sound discrimination tasks were within the range of those reported by previous studies that used 2AFC designs (Lee and Green, 1994; Moore, 1973; Moore *et al*., 1985; Wier *et al*., 1977). We further show that, unlike humans, ferrets’ performance differed drastically across these two task types. Ferrets could detect a 3% change in the reference F0 during the go/no-go task, but on average required a 73% F0 difference to reach criterion on the 2AFC task. Therefore, the threshold differences across these tasks appear to be small for human listeners, but marked for our animal model.

What does this result mean for the comparisons of human and animal discrimination thresholds that are so often made by previous studies? It suggests that differences between human 2AFC and animal go/no-go discrimination performance may well approximate differences in the ability of these species to detect a change in a stimulus parameter (i.e. performance of humans and animals on a common go/no-go task). However, such a comparison may grossly underestimate the differences between humans and animals in judging the direction of a pitch change. It is better practice instead to test all species under study using tasks that are as similar as possible in design and perceptual requirements, and with low cognitive loads.

### 4.2 Possible reasons for ferrets’ performance differences between the 2 tasks

The poor performance of ferrets on the 2AFC task, compared to the go/no-go task, may result from task differences in cognitive demands (that is, the 2AFC may simply be a more procedurally difficult to learn), perceptual demands (labeling pitch shifts may be more perceptually challenging than detecting a pitch change), or motivation to perform the task. We will consider each of these possibilities in turn.

#### 4.2.1 Cognitive demands

In our 2AFC task, the ferrets must label a pitch change as “high” or “low” on each trial by choosing between one of two possible responses. To successfully complete this task, they must attend to the stimuli, identify the type of pitch change (“increase” versus “decrease”), map this identity onto a choice of behavioral responses (here, “left” versus “right” spout), and then produce the appropriate motor activity to carry out the chosen behavioral response (for example, move to the left spout and trigger it). By comparison, a go/no-go change detection task requires the animals to attend to the stimuli, detect a pitch change, and then make the single possible behavioral response (here, release the spout). Having to map the learned perceptual categories of high and low pitch systematically onto different behavioral responses creates a cognitive task demand, which could limit ferrets’ performance of the 2AFC task independent of their perceptual acuity. Simulated models have suggested that inattention can affect performance on 2AFC tasks more severely than on go/no-go tasks (Green, 1995), so it is possible that inattention in ferrets might have contributed to the task-dependency we observed.

It is widely held that that go/no-go tasks are less cognitively demanding than their 2AFC counterparts, and this view is supported by studies showing that animals learn to make sound quality discriminations on the former task much more quickly than the latter (Lawicka, 1964; Burdick, 1980). In contrast, we found no difference in the number of sessions or trials required by ferrets to learn these two pitch discrimination tasks. The number of trials required for task acquisition depend, however, on the stimulus comparisons required as well as the training procedures, apparatus design and type of reinforcement used. These factors vary considerably from study to study. Consequentially, acquisition times are not ideally suited for comparing the cognitive demands of the final testing task itself. Therefore, while our tasks were well matched in acquisition times, this does necessarily imply that the cognitive demands of the final go/no-go and 2AFC tasks were matched in cognitive load.

Perhaps a more relevant observation for this argument is that the ferrets performed the 2AFC task correctly on 80-100% of trials when the pitch discrimination was perceptually easiest (Fig. 2). This indicates that the animals were capable of reliably associating the direction of pitch shift with the appropriate response type when perceptual uncertainty was low.

#### 4.2.2 Perceptual demands

Given the above observations, we think it is unlikely that differences in cognitive demands alone can account for ferrets’ higher thresholds on the 2AFC task. Differences in the perceptual demands of the two tasks may also play a role. On the go/no-go task, it is sufficient to detect that the pitches of two consecutive stimuli are not the same. In the 2AFC task, the animals must identify whether the second pitch is "higher" or "lower" than the first. Identifying the direction of pitch changes requires different computations, and may involve different neurophysiological processes, than merely detecting a change. This is supported by previous studies showing that most human listeners perform marginally better when asked whether two tones have the same pitch than when required to report the direction of a pitch change between the two tones (Creelman and Macmillan, 1979; Neuhoff *et al*., 2002; Semal and Demany, 2006). By contrast, we found that humans did not produce significantly different F0 thresholds on a pitch change detection task and 2AFC pitch direction judgment task, although our go/no-go task was procedurally very different from the AFC same/different tasks employed by the above authors. This again emphasizes the dependence of pitch discrimination thresholds on task design.

The prior experience of the subjects is another factor that needs to be considered since Semal and Demany (2006) found that among subjects with extensive musical experience, who are likely to be used to making pitch comparisons between successive musical notes, up/down pitch comparison thresholds are subtly smaller than those on a same/different pitch task.

Some evidence for an anatomical segregation of the cortical substrates necessary for pitch change detection and pitch comparisons can be found in lesion studies. Johnsrude *et al*. (2000) showed that neurological patients with damage to right Heschl’s gyrus could detect changes in the frequency of tones as well as healthy controls, but were impaired in reporting whether these frequency changes were rising or falling. Similarly, Elliott and Trahiotis (1972) reported that auditory cortical lesions in animals disrupt performance on a frequency recognition task more so than a frequency change detection task.

Demany and Ramos (2005) have suggested that the human auditory system may contain neurons that act as dedicated pitch shift detectors to support up/down pitch judgments, and, if so, animals’ relatively large pitch discrimination thresholds might indicate that they lack such neurons. We have previously shown that ferrets tend to solve the 2AFC pitch direction discrimination task described here by labeling the absolute pitch of the target, whereas human subjects report that their judgments are instead based on the relative pitch of the target with respect to the preceding reference sound (Walker *et al*., 2009). Yin *et al*. (2010) have shown that, when the task requires it, ferrets can be trained to judge the relative frequencies of pure tones. It therefore seems that ferrets, like birds (Dooling *et al*., 1987; Hulse and Cynx, 1985; Page *et al*., 1989) and non-human primates (Brosch et al., 2004; D’Amato, 1988), can make relative frequency judgments but prefer strategies based on labeling absolute pitch (but see also Elliott *et al*., 1971).

It is also remains possible that anatomically overlapping but physiologically distinct neural subpopulations are required for pitch change detection and pitch shift comparisons. Across a variety of animal species, the firing rates of a distributed subset of auditory cortical neurons has been shown to be monotonically modulated by the repetition rate of complex sounds (Eggermont, 1991; Steinschneider *et al*., 1998; Bendor and Wang, 2010; Bizley *et al*., 2010). Bendor and Wang (2010) have suggested that these responses may provide the neural signal for pitch comparisons, while a specialized class of more anatomically restricted auditory cortical neurons may be used to extract a sound’s absolute pitch (Bendor and Wang, 2005). The extent to which any of these observed neural responses contributes to animals’ perception of pitch is currently unclear, but simultaneous measurements of physiology and behavior should help to resolve whether a neural substrate exists for the differences in performance we observe here on 2AFC and go/no-go tasks.

#### 4.2.3 Motivation effects

Our ferrets were given water and time-outs to reinforce their behavior on the 2 tasks, while human subjects were reinforced with visual feedback and the same time-outs. It is impossible to say whether the motivation to respond accurately was the same in each species. However, as the rewards and time-outs were matched within species, motivational effects cannot account for differences in ferrets’ thresholds between the go/no-go and 2AFC tasks. Previous animal psychoacoustical studies have used positive reinforcement (food or water rewards), punishment (usually electric shock), or a combination of the two to train the behavioral task. The frequency difference thresholds measured within a species do not differ markedly across these reinforcement regimes (compare Massopust *et al*., 1971 with Sinnott *et al*., 1987 or Stebbins, 1973; Kelly, 1979 with Talwar & Gerstein, 1998). Positive reinforcement offers advantages, however, in terms of animal welfare and the brain mechanisms involved (David *et al*., 2012; Masterton, 1997).

## 5. CONCLUSIONS

Experiments that combine behavioral training with electrophysiological recordings and neural deactivation in animals hold promise for elucidating the neural substrates of pitch perception. However, before embarking on such studies, it is important to appreciate which perceptual abilities are assessed via behavioral tasks. This study has shown that two classic 2AFC and go/no-go task designs result in pitch discrimination thresholds that differ substantially in magnitude in one species (ferrets), but not in another (humans). The extent to which the observed performance differences across 2AFC and go/no-go tasks are due to cognitive or perceptual factors remains unclear, but there are reasons to expect that both might contribute. These results emphasize the particular sensitivity of animal psychophysical thresholds to task design, and thus the importance of careful behavioral measurements of perceptual decisions that are under neurophysiological investigation in an animal species.

## ACKNOWLEDGEMENTS

The authors thank Sheelah Hart-Schnupp, Chui Tsang and Sarah Wynbourne for assistance with ferret behavioral training.

## Footnotes

1 The F0 discrimination performance of these 2 animals (denoted with symbols “+” and “x”) was described in Walker et al. (2009), but are not included in the present study because they were not tested with a reference of 400 Hz.

